# Genomic surveillance of *Bacillus cereus sensu lato* strains isolated from meat and poultry products in South Africa enables inter- and intra-national surveillance and source tracking

**DOI:** 10.1101/2022.01.18.476746

**Authors:** Laura M. Carroll, Rian Pierneef, Aletta Mathole, Abimbola Atanda, Itumeleng Matle

## Abstract

Members of the *Bacillus cereus sensu lato* (*s.l.*) species complex, also known as the *B. cereus* group, vary in their ability to cause illness, but are frequently isolated from foods, including meat products; however, food safety surveillance efforts that employ whole-genome sequencing (WGS) often neglect these potential pathogens. Here, WGS was used to characterize *B. cereus s.l*. strains (*n* = 25) isolated during surveillance of meat products in South Africa. Strains were collected from beef, poultry, and mixed meat products obtained from (i) retail outlets, processing plants, and butcheries across six South African provinces (*n* = 15, 7, and 1, respectively), and (ii) imports in cold stores (*n* = 2). Strains were assigned to *panC* Groups IV, III, II, and V (*n* = 18, 5, 1, and 1, respectively) and spanned multiple genomospecies, regardless of the taxonomy used. All strains possessed diarrheal toxin-encoding genes, while one sequence type 26 (ST26) strain possessed cereulide (emetic toxin) synthetase-encoding genes. No strains harbored anthrax toxin- or capsule-encoding genes. The 25 strains were partitioned into 15 lineages via *in silico* seven-gene multi-locus sequence typing (MLST), six of which contained multiple strains sequenced in this study, which were identical or nearly identical at the whole-genome scale. Five MLST lineages contained (nearly) identical genomes collected from two or three South African provinces; one MLST lineage contained nearly identical genomes from two countries (South Africa and the Netherlands), indicating that *B. cereus s.l*. can spread intra- and inter-nationally via foodstuffs.

**Importance:** Nation-wide foodborne pathogen surveillance programs that employ high-resolution genomic methods have been shown to provide vast public health and economic benefits. However, *B. cereus s.l*. are often overlooked during large-scale, routine WGS efforts. Thus, to our knowledge, no studies to date have evaluated the potential utility of WGS for *B. cereus s.l*. surveillance and source tracking in foodstuffs. In this proof-of-concept study, we applied WGS to *B. cereus s.l*. strains collected via South Africa’s national surveillance program of domestic and imported meat products, and we provide strong evidence that *B. cereus s.l*. can be disseminated intra- and inter-nationally via the agro-food supply chain. Our results showcase that WGS can be used for source tracking of *B. cereus s.l*. in foods, although future WGS and isolate metadata collection efforts are needed to ensure that *B. cereus s.l*. surveillance initiatives are on par with those of other foodborne pathogens.

## INTRODUCTION

*Bacillus cereus sensu lato (s.l.*), also known as the *B. cereus* group, is a complex of closely related, Gram-positive, spore-forming species, which are widespread throughout the environment (1). While some members of *B. cereus s.l*. have important industrial applications or roles (e.g., as biocontrol agents in agricultural settings, as food spoilage organisms) (2–6), others are capable of causing illnesses or death in humans and/or animals (1, 7–9). Illnesses caused by members of *B. cereus s.l*. can range in severity from mild to severe/fatal and include anthrax and anthrax-like illness (8, 10, 11), foodborne emetic intoxication (1, 7, 12–15), foodborne diarrheal toxicoinfection (1, 7, 14, 15), and non-gastrointestinal infections (16, 17). As a foodborne pathogen, “*B. cereus*” is estimated to be responsible for more than 256,000 illnesses globally each year (18), although this is likely an underestimate, due to the relatively mild and self-limiting nature of the symptoms that often accompany foodborne illness caused by *B. cereus s.l*. (1).

Food safety surveillance efforts around the world have identified *B. cereus s.l*. strains in a wide variety of foodstuffs (1, 7), including raw intact, processed, and ready-to-eat (RTE) meat and poultry products (19–27). In South Africa specifically, previous surveillance efforts have identified *B. cereus s.l*. in (i) retail meats sold at supermarkets in the Pretoria area (i.e., Vienna sausages, salami, and poultry) (27) and (ii) biltong (a spiced intermediate moisture RTE meat product) sold at supermarkets, stalls, kiosks, and butcheries in Bloemfontein, Free State (28). Most recently, in a study of over two thousand meat product samples collected from butcheries, processing plants, abattoirs, and retail outlets across all nine South African provinces, *B. cereus s.l*. was present in 4.5 and 2.7% of domestic and imported meat products, respectively (26).

Ongoing surveillance efforts in South Africa have indicated that meat and poultry products can harbor *B. cereus s.l*. and may pose a potential food safety risk to South African consumers. However, it is unclear which *B. cereus s.l*. lineages are present in South African meat and poultry products on a genomic scale. Here, we used whole-genome sequencing (WGS) to characterize 25 *B. cereus s.l*. strains isolated from raw intact, processed, and RTE meat and poultry products collected from processing plants, butcheries, and retail outlets across South Africa, as well as imported meat products. By comparing South African strains sequenced here to all publicly available *B. cereus s.l*. genomes (*n* = 2,887 total genomes), we identified multiple *B. cereus s.l*. species present among South African meat and poultry products, and we detected multiple potential inter-national and inter-provincial *B. cereus s.l*. dissemination events. Overall, our study serves as the first genome-scale study of South African *B. cereus s.l*. in foodstuffs and showcases the utility of WGS for *B. cereus s.l*. surveillance and source tracking.

## RESULTS

### *B. cereus s.l*. are present among domestic and imported meat and poultry products in South Africa

A total of 25 *B. cereus s.l*. strains were isolated from meat and poultry products, which had been collected across South Africa in 2015 and 2016 (Figure 1, Table 1, and Supplemental Table S1). Overall, 19 strains (76.0%) originated from beef products, including: beef wors, a processed South African sausage (*n* = 7; 28.0%); beef biltong, a South African spiced intermediate moisture RTE meat product (*n* = 5; 20.0%); processed beef mince (*n* = 3; 12.0%); RTE beef sausage emulsion (*n* = 2; 8.0%); and processed beef patties (*n* = 2; 8.0%, Table 1 and Supplemental Table S1). Five strains (20.0%) were isolated from poultry products, including raw chicken thighs (*n* = 2; 8.0%), raw chicken quarter-legs (*n* = 2; 8.0%), and a frankfurter (*n* = 1; 4.0%, Table 1 and Supplemental Table S1). One strain (4.0%) was isolated from wors, which had been made from a mix of beef, pork, and lamb (Table 1 and Supplemental Table S1).

**Figure 1.**
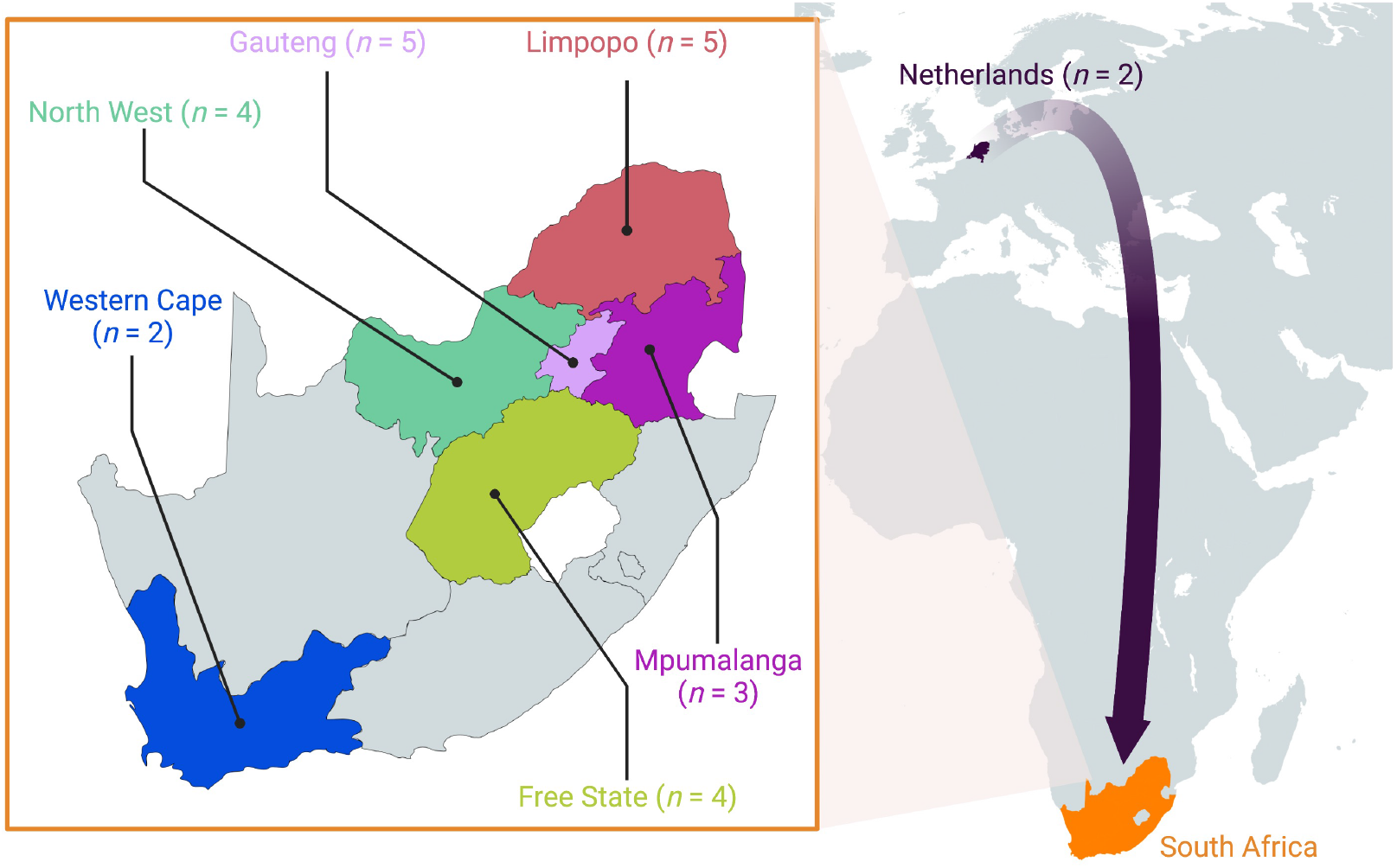
Geographic origins of *B. cereus s.l*. strains sequenced in this study (*n* = 25).

**Table 1.** *B. cereus s.l*. strains sequenced in this study (*n* = 25).

| Strain | Year<sup>a</sup> | Meat Sample Type | Meat Category<sup>b</sup> | Food Animal | Establishment | Province<sup>c</sup> | *panC*<sup>d</sup> Group | MLST ST<sup>e</sup> | GTDB Species<sup>f</sup> | 2020 GSB Species<sup>g</sup> |
| --- | --- | --- | --- | --- | --- | --- | --- | --- | --- | --- |
| S57 | 2015 | Beef Biltong | RTE | Beef | Retail Outlets | Free State | II | 794 | *wiedmannii* | *mosaicus* |
| S58 | 2015 | Chicken Thigh | Raw Intact | Poultry | Retail Outlets | Mpumalanga | III | NA | *anthracis*<sup>h</sup> | *mosaicus* |
| S59 | 2016 | Chicken 1/4 Leg | Raw Intact | Poultry | Cold Store | Import (NL) | III | 2413 | *paranthracis* | *mosaicus* |
| S62 | 2015 | Chicken Thigh | Raw Intact | Poultry | Retail Outlets | Mpumalanga | III | 2413 | *paranthracis* | *mosaicus* |
| S64 | 2016 | Sausage Emulsion | RTE | Beef | Retail Outlets | Western Cape | III | NA | *anthracis*<sup>h</sup> | *mosaicus* |
| S66 | 2015 | Beef Mince | Processed | Beef | Processing Plant | Mpumalanga | III | 26 | *paranthracis* | *mosaicus*<br>subsp. *cereus* |
| S51 | 2015 | Beef Mince | Processed | Beef | Retail Outlets | Limpopo | IV | 1578 | *thuringiensis* | *cereus* s.s. |
| S53 | 2015 | Beef Patties | Processed | Beef | Retail Outlets | Gauteng | IV | 2668 | *thuringiensis*_S | *cereus* s.s. |
| S55 | 2015 | Beef Wors | Processed | Beef | Processing Plant | Limpopo | IV | 177 | *cereus* | *cereus* s.s. |
| S56 | 2015 | Beef Wors | Processed | Beef | Processing Plant | North West | IV | 2721 | *bombysepticus* | *cereus* s.s. |
| S63 | 2015 | Beef Biltong | Processed | Beef | Retail Outlets | Free State | IV | 2668 | *thuringiensis*_S | *cereus* s.s. |
| S65 | 2015 | Beef Wors | Processed | Beef | Processing Plant | Limpopo | IV | NA | *bombysepticus* | *cereus* s.s. |
| S67 | 2016 | Chicken 1/4 Leg | Raw Intact | Poultry | Cold Store | Import (NL) | IV | NA | *bombysepticus* | *cereus* s.s. |
| S70 | 2015 | Beef Mince | Processed | Beef | Retail Outlets | Limpopo | IV | 2668 | *thuringiensis*_S | *cereus* s.s. |
| S77 | 2015 | Beef Wors | Processed | Beef | Retail Outlets | Gauteng | IV | 24 | *cereus* | *cereus* s.s. |
| S78 | 2015 | Beef Biltong | RTE | Beef | Retail Outlets | Gauteng | IV | 1697 | *thuringiensis*_S | *cereus* s.s. |
| S79 | 2015 | Beef Biltong | RTE | Beef | Retail Outlets | North West | IV | 2721 | *bombysepticus* | *cereus* s.s. |
| S80 | 2015 | Beef Wors | Processed | Beef | Processing Plant | North West | IV | 177 | *cereus* | *cereus* s.s. |
| S81 | 2016 | Beef Wors | Processed | Beef | Processing Plant | North West | IV | NA | *cereus* | *cereus* s.s. |
| New_S84 | 2015 | Beef Patties | Processed | Beef | Retail Outlets | Free State | IV | 2721 | *bombysepticus* | *cereus* s.s. |
| S85 | 2015 | Beef Wors | Processed | Beef | Retail Outlets | Gauteng | IV | NA | *bombysepticus* | *cereus* s.s. |
| S86 | 2015 | Sausage Emulsion | RTE | Beef | Retail Outlets | Western Cape | IV | 2289 | *cereus* | *cereus* s.s. |
| S87 | 2015 | Frankfurter | Raw Intact | Poultry | Processing Plant | Free State | IV | NA | *bombysepticus* | *cereus* s.s. |
| S88 | 2015 | Beef Biltong | RTE | Beef | Retail Outlets | Limpopo | IV | NA | *cereus* | *cereus* s.s. |
| S72 | 2015 | Beef-Pork-Lamb Wors | Processed | Mixed | Butchery | Gauteng | V | 223 | *toyonensis* | *toyonensis* |

<sup>a</sup>Year of isolation
<sup>b</sup>RTE, Ready-to-Eat
<sup>c</sup>Two strains were isolated from meat products imported from the Netherlands (NL)
<sup>d</sup>*panC* phylogenetic group assigned using BTyper3
<sup>e</sup>Sequence type (ST) assigned using the PubMLST seven-gene multi-locus sequence typing (MLST) scheme for *B. cereus* and BTyper3
<sup>f</sup>Genome Taxonomy Database (GTDB) species assigned using GTDB-Tk
<sup>g</sup>Species and subspecies (where applicable), assigned using the 2020 Genomospecies-Subspecies-Biovar (GSB) nomenclatural framework for *B. cereus s.l.* and BTyper3 (see Table 2 for predicted biovar/phenotypic information)
<sup>h</sup>Despite GTDB assigning a species label of “*B. anthracis*”, these strains cannot cause anthrax illness, nor do they belong to the classic “*B. anthracis*” lineage most commonly responsible for anthrax illness (34)

The majority of strains (23 of 25, 92.0%) were obtained from domestic meat and poultry products acquired across six South African provinces (*n* = 5, 5, 4, 4, 3, and 2 strains from Gauteng, Limpopo, Free State, North West, Mpumalanga, and Western Cape, respectively; Figure 1, Table 1, and Supplemental Table S1). The remaining two strains (8.0%) were isolated from raw, intact chicken quarter-legs, which had been imported into South Africa from the Netherlands (Figure 1, Table 1, and Supplemental Table S1). Fifteen of the 25 *B. cereus s.l*. strains (60.0%) were isolated from meat or poultry products acquired from retail outlets, while seven (28.0%) were acquired from meat or poultry products obtained from processing plants (Table 1 and Supplemental Table S1). Two strains (from chicken quarter-legs imported from the Netherlands; 8.0%) were derived from poultry in cold stores, while one strain (4.0%) was isolated from the mixed beef-pork-lamb wors acquired from a butchery (Table 1 and Supplemental Table S1).

### Multiple species are present among *B. cereus s.l*. from South African meat and poultry products

Species-level taxonomic classification of *B. cereus s.l*. is notoriously challenging (1, 29, 30); to avoid taxonomic ambiguities and maximize interpretability, we applied multiple taxonomic assignment and sequence typing methods to the 25 strains sequenced here (Table 1). One such sequence typing framework relies on the pantoate-β-alanine ligase gene (*panC*) to assign *B. cereus s.l*. strains to one of seven or more major phylogenetic groups, which have been proposed to conceptually serve as “species” (31, 32). Using the adjusted eight-group *panC* typing approach implemented in BTyper3 (33), the 25 *B. cereus s.l*. strains sequenced here encompassed four *panC* phylogenetic groups (i.e., “species”; Figure 2 and Table 1). The majority (*n* = 18 of 25, 72.0%) of the strains sequenced here were assigned to *panC* Group IV (Figure 2 and Table 1). The remaining strains were assigned to *panC* Groups III, II, and V (*n* = 5, 1, and 1 strains, representing 20.0%, 4.0%, and 4.0% of isolates sequenced here, respectively; Figure 2 and Table 1).

**Figure 2.**
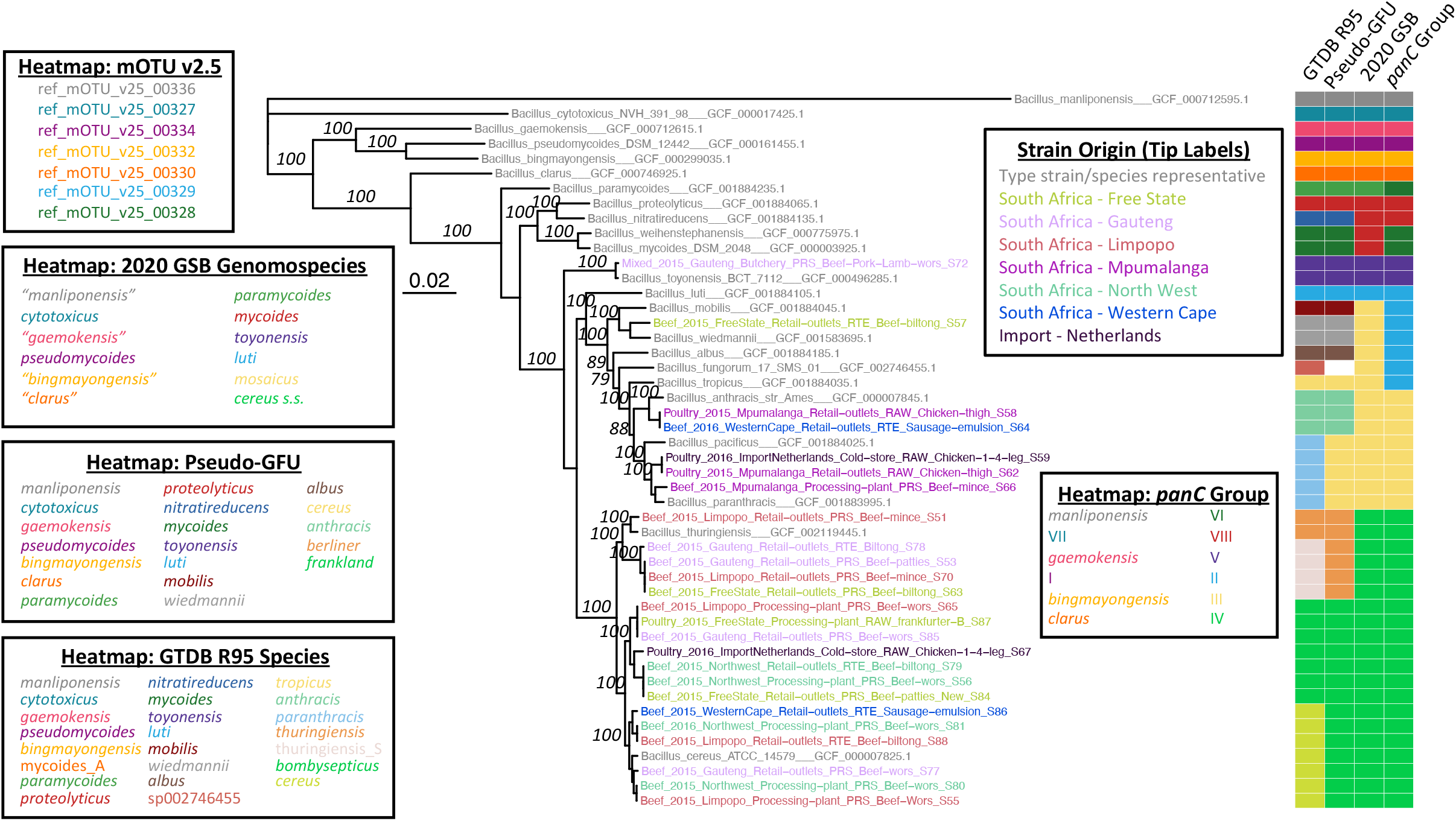
Maximum likelihood (ML) phylogeny constructed using amino acid sequences derived from the 25 *B. cereus s.l*. isolate genomes sequenced in this study (tip labels colored by geographic origin), plus type strain/species representative genomes of 23 published and effective *B. cereus s.l*. species (gray tip labels). The heatmap to the right of the phylogeny showcases species assignments obtained within the following taxonomic frameworks (from left to right): (i) Genome Taxonomy Database (GTDB) Release 05-RS95 and GTDB-Tk (GTDB R95); (ii) pseudo-gene flow units (GFUs) assigned using BTyper3 (Pseudo-GFU); (iii) genomospecies of the 2020 standardized *B. cereus s.l*. genomospecies/subspecies/biovar (GSB) framework and BTyper3 (2020 GSB); (iv)*panC* Group (I-VIII), assigned using BTyper3 (*panC* Group). The phylogeny was constructed using IQ-TREE, using core orthologs identified among all genomes via OrthoFinder as input. Branch lengths are reported in substitutions per site. Branch labels correspond to branch support percentages obtained using 1,000 replicates of the ultrafast bootstrap approximation (selected for readability). The type strain genome of effective *B. cereus s.l*. species “*B. manliponensis*” (the most distant recognized member of *B. cereus s.l.*) was used to root the phylogeny. Heatmap legends for all five taxonomies are colored by their order of appearance in the heatmap, from top to bottom; white heatmap cells denote genomes that could not be assigned to a taxonomic unit within a given taxonomic framework.

Using the Genome Taxonomy Database (GTDB) taxonomy, the 25 *B. cereus s.l*. strains sequenced here encompassed eight genomospecies (Figure 2 and Table 1). The 18 *panC* Group IV strains sequenced here encompassed four GTDB genomospecies, while the five *panC* Group III strains spanned two GTDB genomospecies (Figure 2 and Table 1). The *panC* Group II and Group V strains (*n* = 1 each) were each assigned to separate GTDB genomospecies (Figure 2 and Table 1).

Using a standardized genomospecies-subspecies-biovar (GSB) nomenclatural framework proposed for *B. cereus s.l*. in 2020 (34) (referred to hereafter as the “2020 GSB” framework), the 25 strains sequenced here encompassed three genomospecies (Figure 2 and Tables 1 and 2). All 18 *panC* Group IV strains were assigned to the *B. cereus sensu stricto* (*s.s*.) genomospecies (Figure 2 and Tables 1 and 2); genes encoding insecticidal toxins (referred to hereafter as “Bt toxin-encoding genes”) were detected within all Group IV *B. cereus s.s*. genomes (using BtToxin_scanner2’s “old” gene detection approach), meaning that these 18 strains were predicted to belong to the Thuringiensis biovar (i.e., *B. cereus s.s*. bv. Thuringiensis; Table 2). All five *panC* Group III and the single *panC* Group II strain(s) were assigned to genomospecies *B. mosaicus* within the 2020 GSB framework (Figure 2 and Tables 1 and 2). One *panC* Group III *B. mosaicus* strain (i.e., strain S66) was assigned to PubMLST sequence type 26 (ST26) and possessed cereulide (emetic toxin) synthetase-encoding *cesABCD* and was thus assigned to the *cereus* subspecies and biovar Emeticus (i.e., *B. mosaicus* subsp. *cereus* bv. Emeticus; Table 2). The lone *panC* Group V strain sequenced here was assigned to the *B. toyonensis* genomospecies; Bt toxin-encoding genes were detected in this genome (via BtToxin_scanner2’s “old” gene detection approach), and thus this strain was predicted to belong to biovar Thuringiensis (i.e., *B. toyonensis* bv. Thuringiensis; Figure 2 and Tables 1 and 2).

**Table 2.**
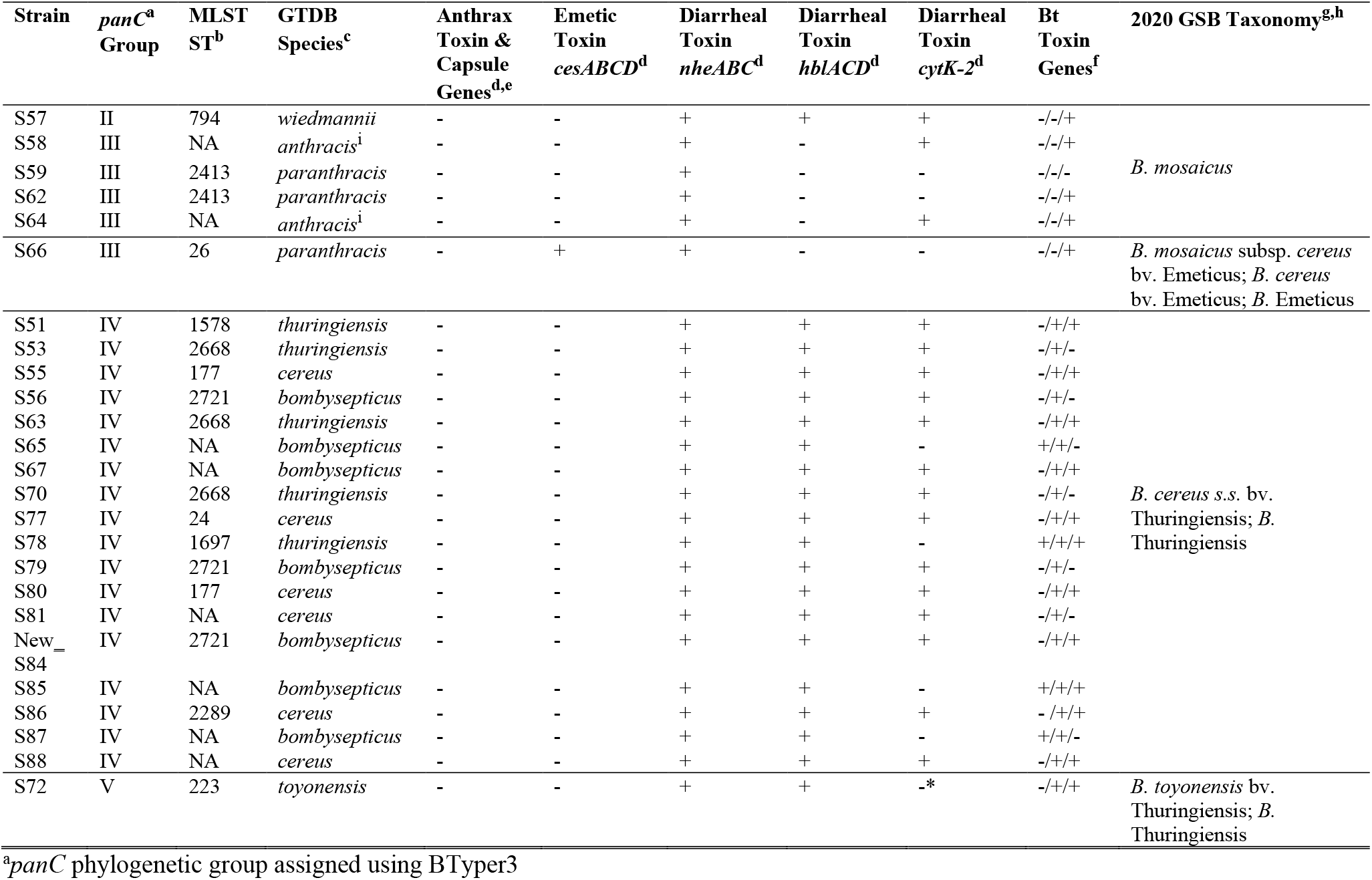
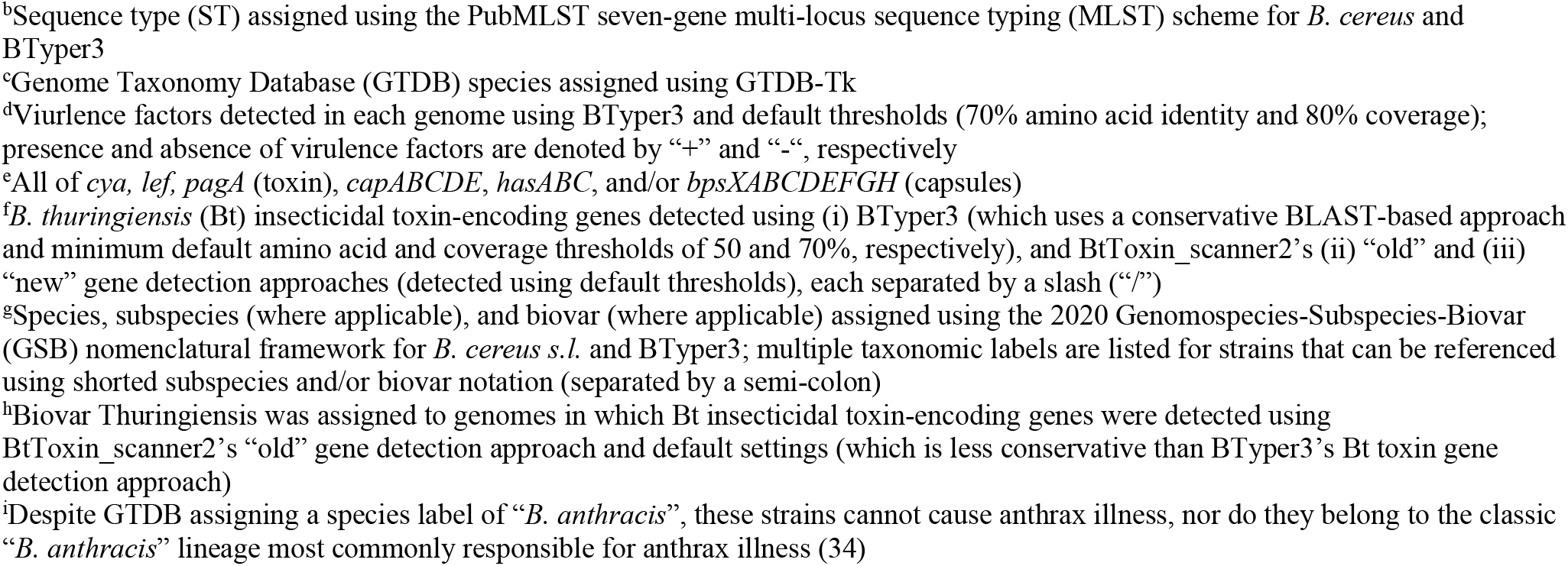
Predicted phenotypes of *B. cereus s.l*. strains sequenced in this study (*n =* 25).

| Strain | *panC*<sup>a</sup><br>Group | MLST<br>ST<sup>b</sup> | GTDB<br>Species<sup>c</sup> | Anthrax<br>Toxin &<br>Capsule<br>Genes<sup>d,e</sup> | Emetic<br>Toxin<br>*cesABCD*<sup>d</sup> | Diarrheal<br>Toxin<br>*nheABC*<sup>d</sup> | Diarrheal<br>Toxin<br>*hblACD*<sup>d</sup> | Diarrheal<br>Toxin<br>*cytK-2*<sup>d</sup> | Bt<br>Toxin<br>Genes<sup>f</sup> | 2020 GSB Taxonomy<sup>g,h</sup> |
| --- | --- | --- | --- | --- | --- | --- | --- | --- | --- | --- |
| S57 | II | 794 | *wiedmannii* | - | - | + | + | + | -/-/+ | *B. mosaicus* |
| S58 | III | NA | *anthracis*<sup>i</sup> | - | - | + | - | + | -/-/+ |  |
| S59 | III | 2413 | *paranthracis* | - | - | + | - | - | -/-/- |  |
| S62 | III | 2413 | *paranthracis* | - | - | + | - | - | -/-/+ |  |
| S64 | III | NA | *anthracis*<sup>i</sup> | - | - | + | - | + | -/-/+ |  |
| S66 | III | 26 | *paranthracis* | - | + | + | - | - | -/-/+ | *B. mosaicus* subsp. *cereus*<br>bv. Emeticus; *B. cereus*<br>bv. Emeticus; *B. Emeticus* |
| S51 | IV | 1578 | *thuringiensis* | - | - | + | + | + | -/+/+ | *B. cereus s.s.* bv.<br>Thuringiensis; *B.*<br>Thuringiensis |
| S53 | IV | 2668 | *thuringiensis* | - | - | + | + | + | -/+/- |  |
| S55 | IV | 177 | *cereus* | - | - | + | + | + | -/+/+ |  |
| S56 | IV | 2721 | *bombysepticus* | - | - | + | + | + | -/+/- |  |
| S63 | IV | 2668 | *thuringiensis* | - | - | + | + | + | -/+/+ |  |
| S65 | IV | NA | *bombysepticus* | - | - | + | + | - | +/+/- |  |
| S67 | IV | NA | *bombysepticus* | - | - | + | + | + | -/+/+ |  |
| S70 | IV | 2668 | *thuringiensis* | - | - | + | + | + | -/+/- |  |
| S77 | IV | 24 | *cereus* | - | - | + | + | + | -/+/+ |  |
| S78 | IV | 1697 | *thuringiensis* | - | - | + | + | - | +/+/+ |  |
| S79 | IV | 2721 | *bombysepticus* | - | - | + | + | + | -/+/- |  |
| S80 | IV | 177 | *cereus* | - | - | + | + | + | -/+/+ |  |
| S81 | IV | NA | *cereus* | - | - | + | + | + | -/+/- |  |
| New_<br>S84 | IV | 2721 | *bombysepticus* | - | - | + | + | + | -/+/+ |  |
| S85 | IV | NA | *bombysepticus* | - | - | + | + | - | +/+/+ |  |
| S86 | IV | 2289 | *cereus* | - | - | + | + | + | - /+ /+ |  |
| S87 | IV | NA | *bombysepticus* | - | - | + | + | - | + /+ /- |  |
| S88 | IV | NA | *cereus* | - | - | + | + | + | - /+ /+ |  |
| S72 | V | 223 | *toyonensis* | - | - | + | + | -* | - /+ /+ | *B. toyonensis* bv.<br>Thuringiensis; *B.*<br>Thuringiensis |

<sup>a</sup>*panC* phylogenetic group assigned using BTyper3
<sup>b</sup>Sequence type (ST) assigned using the PubMLST seven-gene multi-locus sequence typing (MLST) scheme for *B. cereus* and BTyper3
<sup>c</sup>Genome Taxonomy Database (GTDB) species assigned using GTDB-Tk
<sup>d</sup>Virulence factors detected in each genome using BTyper3 and default thresholds (70% amino acid identity and 80% coverage); presence and absence of virulence factors are denoted by “+” and “-“, respectively
<sup>e</sup>All of *cya*, *lef*, *pagA* (toxin), *capABCDE*, *hasABC*, and/or *bpsXABCDEFGH* (capsules)
<sup>f</sup>*B. thuringiensis* (Bt) insecticidal toxin-encoding genes detected using (i) BTyper3 (which uses a conservative BLAST-based approach and minimum default amino acid and coverage thresholds of 50 and 70%, respectively), and BtToxin\_scanner2’s (ii) “old” and (iii) “new” gene detection approaches (detected using default thresholds), each separated by a slash (“/”)
<sup>g</sup>Species, subspecies (where applicable), and biovar (where applicable) assigned using the 2020 Genomospecies-Subspecies-Biovar (GSB) nomenclatural framework for *B. cereus s.l.* and BTyper3; multiple taxonomic labels are listed for strains that can be referenced using shorted subspecies and/or biovar notation (separated by a semi-colon)
<sup>h</sup>Biovar Thuringiensis was assigned to genomes in which Bt insecticidal toxin-encoding genes were detected using BtToxin\_scanner2’s “old” gene detection approach and default settings (which is less conservative than BTyper3’s Bt toxin gene detection approach)
<sup>i</sup>Despite GTDB assigning a species label of “*B. anthracis*”, these strains cannot cause anthrax illness, nor do they belong to the classic “*B. anthracis*” lineage most commonly responsible for anthrax illness (34)

Using a rapid, average nucleotide identity (ANI)-based pseudo-gene flow unit assignment scheme, which attempts to assign *B. cereus s.l*. genomes to taxonomic units that mimic *B. cereus s.l*. “species” previously delineated using recent gene flow (33), the 25 strains sequenced here were assigned to six pseudo-gene flow units (pseudo-GFUs; Figure 2 and Supplemental Table S1). The 18 *panC* Group IV and five *panC* Group III strains sequenced here each spanned two pseudo-GFUs (Figure 2 and Supplemental Table S1). The *panC* Group II and Group V strains (*n* = 1 each) were each assigned to separate pseudo-GFUs, respectively (Figure 2 and Supplemental Table S1).

As mentioned above, considerable phenotypic diversity was predicted among the stains sequenced here, as one strain harbored cereulide synthetase-encoding genes, and 19 strains possessed Bt toxin-encoding genes (detected using BtToxin_scanner2’s “old” gene detection approach; Table 2). No anthrax toxin- or capsule-encoding genes were identified within the genomes of the isolates sequenced here (Table 2).

Overall, regardless of whether the *panC*, GTDB, 2020 GSB, or pseudo-gene flow unit assignment frameworks were used, *B. cereus s.l*. strains isolated from meat and poultry products in South Africa were considerably diverse and represented multiple genomospecies (Figure 2, Tables 1 and 2, and Supplemental Table S1). Additionally, using PubMLST’s seven-gene MLST scheme for *B. cereus*, 17 of 25 strains (68.0%) encompassed 11 STs, with eight strains (32.0%) assigned to unknown STs (Table 1 and Supplemental Table S1). Due to the considerable genomic diversity observed among isolates sequenced here, major lineages represented by strains sequenced in this study are discussed individually in detail below, largely within the context of *panC* Groups and/or MLST STs, as these frameworks are well-established (31, 32, 35, 36) and likely the most interpretable to readers.

### Several *B. cereus s.l*. lineages within *panC* Group IV are distributed across multiple South African provinces

Within *panC* Group IV, the 18 strains sequenced here were partitioned into ten lineages using MLST (referred to hereafter as “MLST-based lineages”; Table 3). Based on (i) the whole-genome phylogeny, (ii) pairwise core single-nucleotide polymorphisms (SNPs) identified within MLST-based lineages, and (iii) ANI values calculated within and between MLST-based lineages, four of ten *panC* Group IV MLST-based lineages contained South African strains sequenced in this study, which were highly similar to at least one other strain at the whole-genome level (Figures 3 and 4 and Table 3). One of these lineages (denoted in Table 3 as Lineage IVA) was composed of three South African strains sequenced in this study (S65, S85, and S87), which were assigned to GTDB’s “*B. bombysepticus*” genomospecies and belonged to an unknown ST (Figures 3 and 4 and Table 3). Despite all three genomes being nearly identical (pairwise core SNP distance = 0, pairwise ANI > 99.99; Figure 3 and Table 3), the three strains were isolated from (i) three different establishments and provinces (a processing plant in Limpopo, a retail outlet in Gauteng, and a processing plant in Free State), and (ii) two different types of meat products (two strains from beef wors and one from a poultry frankfurter; Figure 4 and Table 1).

**Figure 3.**
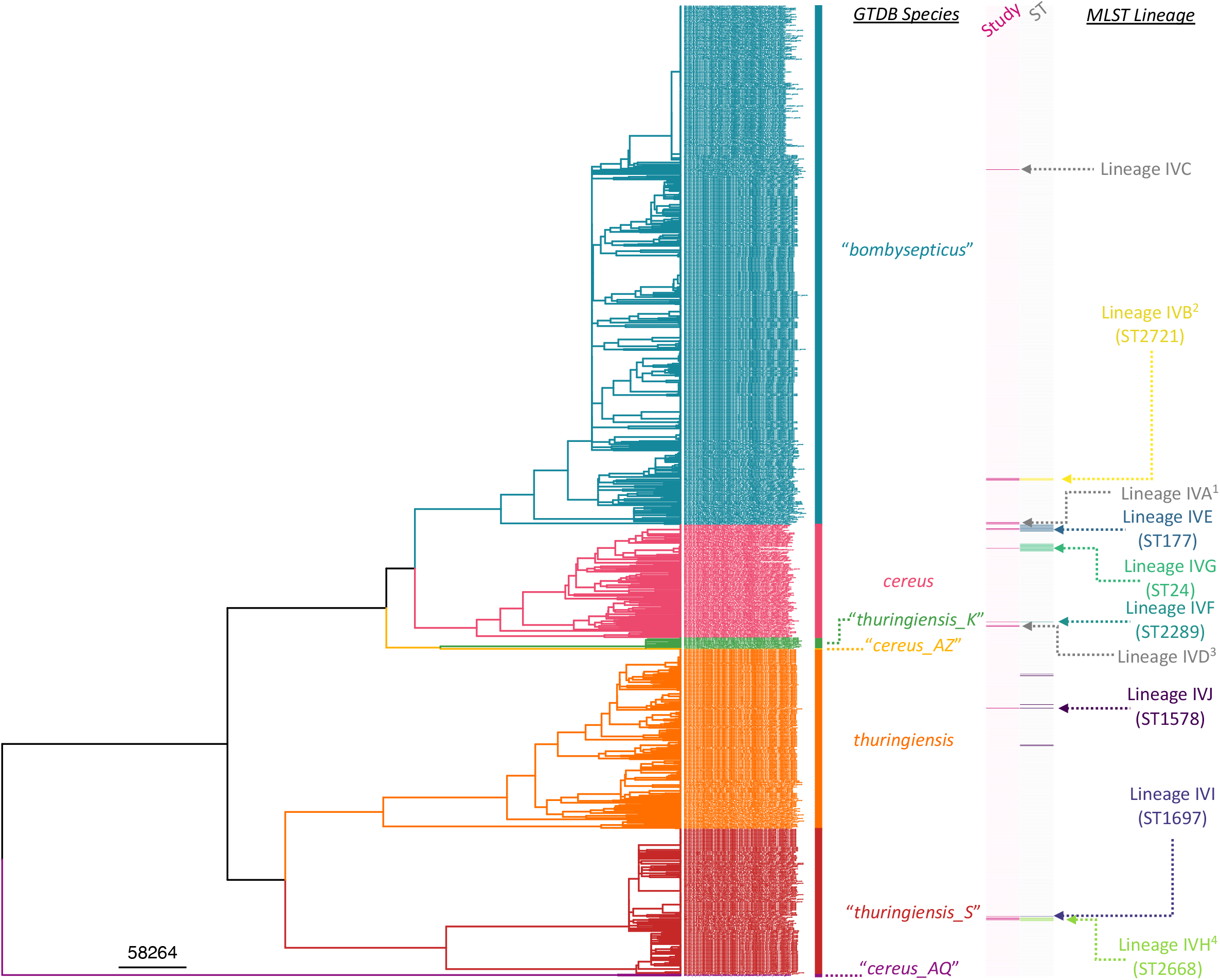
Maximum likelihood (ML) phylogeny constructed using core SNPs identified among orthologous gene clusters of 1,081 *panC* Group IV *B. cereus s.l*. genomes. The phylogeny was rooted using *panC* Group III *B. anthracis* str. Ames Ancestor as an outgroup (NCBI RefSeq Accession GCF_000008445.1; omitted for readability). Tip label colors and clade labels denote species assigned using GTDB-Tk (“GTDB Species”). The heatmap to the right of the phylogeny denotes (i) whether a strain was sequenced in this study (dark pink) or not (light pink; “Study”), and (ii) multi-locus sequence typing (MLST) sequence types (STs) associated with strains sequenced in this study, where applicable (colored), or not (gray; “ST”). MLST lineages discussed in Table 3 are annotated to the right of the heatmap (“MLST Lineage”). MLST lineages with numerical superscripts contain two or more strains sequenced in this study, which were highly similar on a genomic scale; these lineages are depicted in Figure 4. The tree was rooted and time-scaled using LSD2, with branch lengths reported in years.

**Figure 4.**
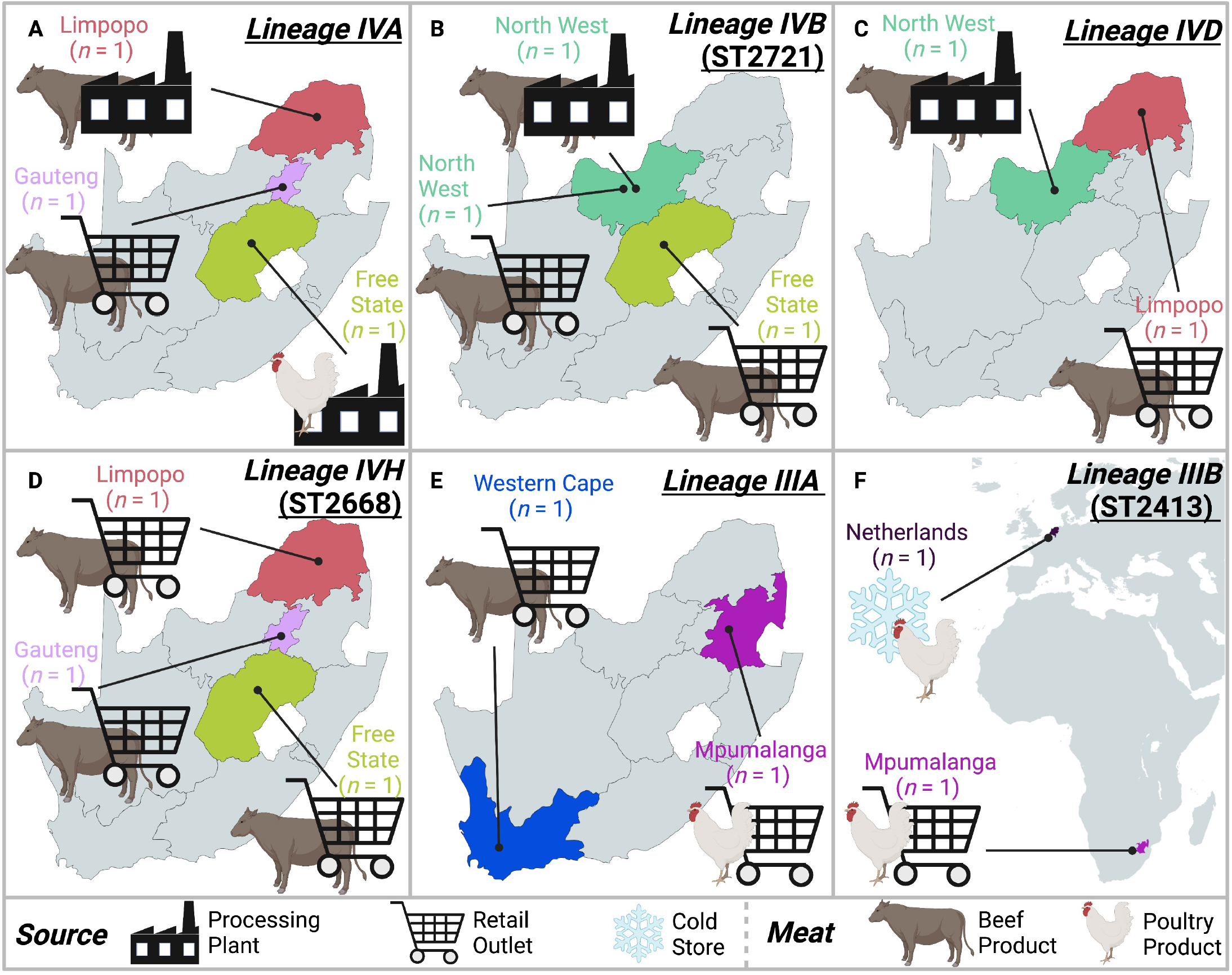
*B. cereus s.l*. multi-locus sequence typing (MLST) lineages that contained two or more strains sequenced in this study, which were identical or nearly identical at the whole-genome scale. Lineage names and sequence types (STs), where applicable, are shown in the top right corner of each panel. Geographic and source origins of each strain are displayed in the respective map.

**Table 3.** Genomic distances within multi-locus sequence typing (MLST) lineages, which contain strains sequenced in this study.

| MLST Lineage<sup>a</sup> | GTDB Species<sup>b</sup> | MLST ST<sup>c</sup> | # of Genomes |  | ANI Range (Mean)<sup>d</sup> | Reference Genome<sup>e</sup> | Pairwise SNP Range within MLST Lineage |  |  |
| --- | --- | --- | --- | --- | --- | --- | --- | --- | --- |
|  |  |  | Total | Study |  |  | Total (Mean)<sup>f</sup> | Within-Study (Mean)<sup>g</sup> | Study-Public (Mean)<sup>h</sup> |
| IVA* | *bombysepticus* | NA | 3 | 3 | 99.99-100.0 (100.0) | S85 | 0-0 (0) | 0-0 (0) | NA |
| IVB* | *bombysepticus* | 2721 | 3 | 3 | 99.99-100.0 (100.0) | S79 | 0-0 (0) | 0-0 (0) | NA |
| IVC | *bombysepticus* | NA | 1 | 1 | NA | NA | NA | NA | NA |
| IVD* | *cereus* | NA | 2 | 2 | 100.0-100.0 (100.0) | S81 | 3 | 3 | NA |
| IVE | *cereus* | 177 | 11 | 2 | 99.70-100.0 (99.92) | S80 | 1-531 (239.2) | 16 | 10-531 (258.11) |
| IVF | *cereus* | 2289 | 1 | 1 | NA | NA | NA | NA | NA |
| IVG | *cereus* | 24<sup>i,j</sup> | 12 | 1 | 99.75-100.0 (99.91) | S77 | 1-1,075 (328.6) | NA | 94-1,054 (290.3) |
| IVH* | *thuringiensis*_S | 2668 | 4 | 3 | 98.90-100.0 (99.49) | S53 | 0-23,098 (11,549) | 0-1 (0.67) | 23,097- 23,098 (23,097) |
| IVI | *thuringiensis*_S | 1697 | 1 | 1 | NA | NA | NA | NA | NA |
| IVJ | *thuringiensis* | 1578<sup>j</sup> | 6 | 1 | 98.49-99.92 (98.81) | S51 | 2,461- 40,845 | NA | 12,642- 36,103 (29,344) |
| IIA | *wiedmannii* | 794 | 3 | 1 | 99.95-100.0 (99.97) | S57 | 31-105 (78.67) | NA | 31-100 (65.50) |
| IIIA* | *anthracis*<sup>k</sup> | NA | 2 | 2 | 99.84-100.0 (99.92) | S64 | 0 | 0 | NA |
| IIIB* | *paranthracis* | 2413 | 2 | 2 | 100.0-100.0 (100.0) | S59 | 1 | 1 | NA |
| IIIC | *paranthracis* | 26 | 77 | 1 | 99.52-100.0 (99.84) | S66 | 0-12,716 (4,797) | NA | 913-11,406 (3,494) |
| VA | *toyonensis* | 223<sup>j</sup> | 41 | 1 | 98.95-99.99 (99.51) | S72 | 14-10,967 (6,600) | NA | 33-10,882 (4,971) |

<sup>a</sup>Lineage identifiers (IDs) pertain to clusters displayed in Figures 3 (Group IV), 5 (Groups II and III), and 6 (Group V); IDs with an asterisk contain two or more strains sequenced in this study, which were highly similar on a genomic scale; NA, not available/applicable
<sup>b</sup>Genome Taxonomy Database (GTDB) species assigned using GTDB-Tk
<sup>c</sup>Sequence type (ST) assigned using the PubMLST seven-gene MLST scheme for *B. cereus* and BTyper3
<sup>d</sup>FastANI average nucleotide identity (ANI) range and mean values between all genomes in the lineage (excludes self-comparisons)
<sup>e</sup>Strain sequenced in this study, which was used as a reference genome for single-nucleotide polymorphism (SNP) calling among all genomes within the lineage via Snippy
<sup>f</sup>Pairwise SNP distances calculated among all genomes within the lineage (excludes self-comparisons)
<sup>g</sup>Pairwise SNP distances calculated among all genomes within the lineage that were sequenced in this study (excludes self- comparisons)
<sup>h</sup>Pairwise SNP distances calculated between genomes sequenced in this study and public genomes within the same lineage (excludes self-comparisons)
<sup>i</sup>For polyphyletic ST24, one outlier ST24 genome was excluded (NCBI RefSeq Accession GCF\_010580595.1)

Similar results were observed for *panC* Group IV ST2721 (Lineage IVB in Table 3), which was also assigned to GTDB’s “*B. bombysepticus*” genomospecies and contained three strains sequenced in this study (S56, S79, and New_S84; Figures 3 and 4 and Table 3). The three ST2721 genomes were nearly identical (pairwise core SNP distance = 0, pairwise ANI > 99.99; Table 3), despite the fact that the strains originated from three different meat products (one from each of beef wors, beef biltong, and processed beef patties) obtained from three different establishments (from a processing plant in North West province, a retail outlet in North West province, and a retail outlet in Free State, respectively; Figure 4 and Table 1).

A third *panC* Group IV lineage of unknown ST, which was assigned to GTDB’s *B. cereus* species (Lineage IVD), contained two isolates sequenced in this study (S81 and S88; Figures 3 and 4 and Table 3). One strain (S81) had been isolated from beef wors in a processing plant in the North West province; the other (S88) from RTE beef biltong in a retail outlet in Limpopo (Figure 4 and Table 1). Both strains were highly similar on a genomic scale (>99.99 ANI) and differed by three core SNPs (Figure 3 and Table 3). For reference, in a previous point source foodborne outbreak caused by *B. cereus s.l*. (37), outbreak isolates could differ by up to seven core SNPs (using the same SNP calling methodology used here) (38).

A fourth *panC* Group IV lineage, ST2668, contained three strains sequenced in this study (S53, S63, and S70), which were assigned to GTDB’s “*B. thuringiensis_S*” genomospecies (Lineage IVH; Figures 3 and 4 and Table 3). The three strains sequenced here differed by, at most, a single core SNP (Table 3), even though all had been isolated from different meat products (processed beef patties, beef biltong, and processed beef mince) obtained from different establishments (i.e., from retail outlets in each of Gauteng, Free State, and Limpopo, respectively; Figure 4 and Table 1). One publicly available genome associated with a *B. cereus s.l*. strain isolated from grass in KarieDeshe, Israel (39) was additionally assigned to ST2668 (NCBI RefSeq Accession GCF_005217245.1); however, this strain was not closely related to the three nearly identical South African ST2668 strains sequenced in this study (Table 3).

The remaining six MLST-based lineages (i.e., Lineages IVC, IVE, IVF, IVG, IVI, and IVJ, corresponding to an unknown ST, ST177, ST2289, ST24, ST1697, and ST1578, respectively; Table 3) contained isolates sequenced in this study, which were not closely related to any other strains at the whole-genome level (Figure 3 and Table 3). Lineages IVC, IVF, and IVI were singleton lineages, which each contained one genome sequenced in this study (S67, S86, and S78, respectively), shared 98.90-99.35 ANI with their closest publicly available neighbors (Figure 3 and Table 3). Two lineages (IVG and IVJ) each contained multiple genomes, but only one genome sequenced in this study (i.e., S77 and S51, respectively; Figure 3 and Table 3); for both lineages, the South African genome sequenced here was not identical to any publicly available genomes. The remaining lineage, Lineage IVE (ST177), contained multiple genomes, as well as multiple genomes sequenced in this study (Figure 3 and Table 3). Notably, the two ST177 strains sequenced in this study (S55 and S80), which were assigned to this lineage, were not identical, and were isolated from beef wors from processing plants in the Limpopo and North West provinces, respectively (Table 1), indicating that WGS can be useful for differentiating closely related *B. cereus s.l*. genomes within STs.

### A *panC* Group III *B. cereus s.l*. lineage with a novel sequence type was detected in beef and poultry products from two South African provinces

Two *panC* Group III *B. cereus s.l*. strains sequenced here were assigned to an unknown ST (S58 and S64; Figure 5 and Table 1). Despite having been isolated from different products (S58 from a raw chicken thigh, S64 from RTE sausage emulsion) from different establishments (retail outlets in Mpumalanga and Western Cape, respectively), the strains were nearly identical (pairwise core SNP distance = 0, pairwise ANI > 99.84; Figures 4 and 5 and Table 3), indicating that these two strains share a common source.

**Figure 5.**
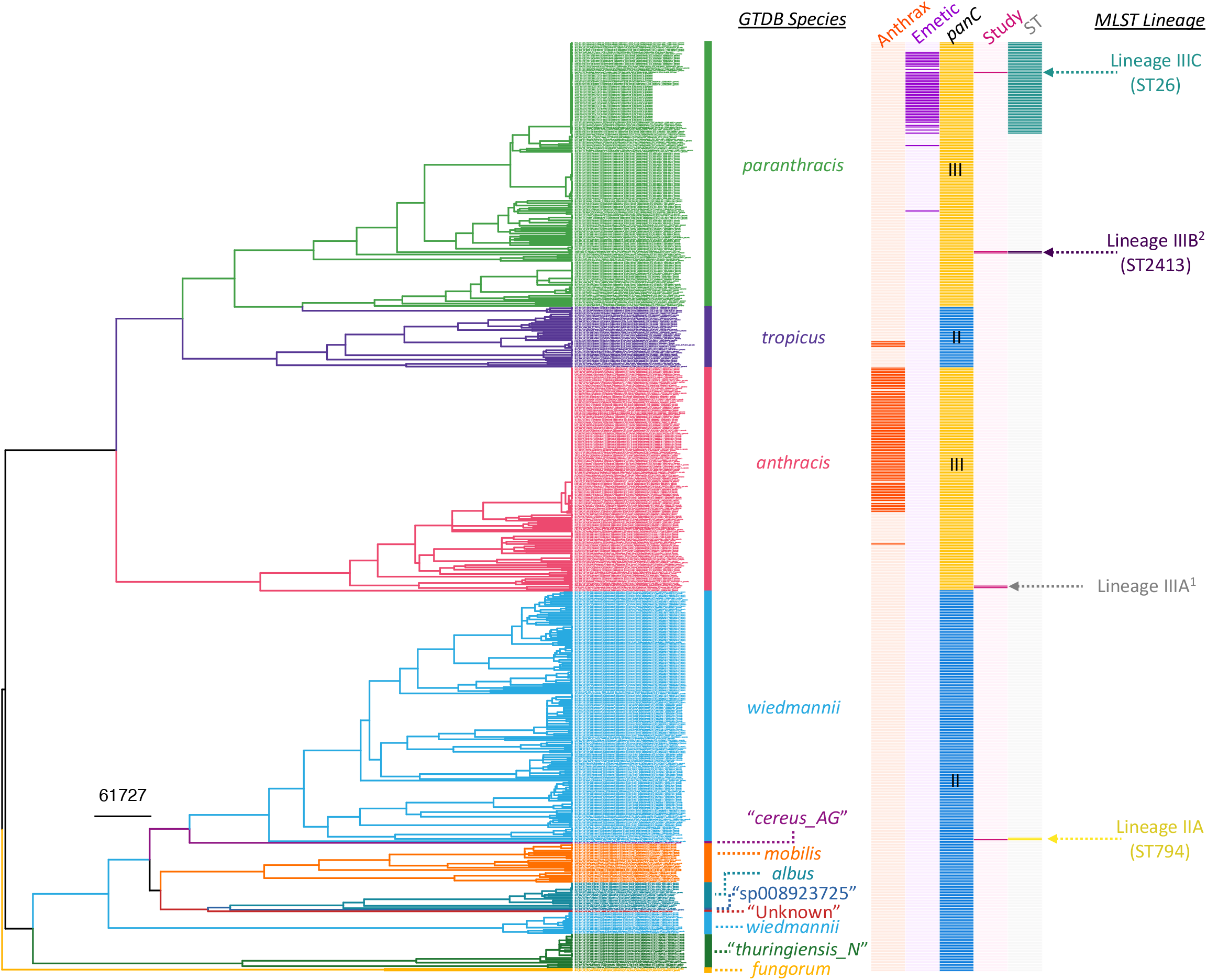
Maximum likelihood phylogeny constructed using core SNPs identified among orthologous gene clusters of 597 *panC* Group II and III *B. cereus s.l*. genomes. The phylogeny was rooted using *panC* Group IV *B. cereus* str. ATCC 14579 as an outgroup (NCBI RefSeq Accession GCF_006094295.1; omitted for readability). Tip label colors and clade labels denote species assigned using GTDB-Tk (“GTDB Species”). The heatmap to the right of the phylogeny denotes (i) whether a strain possesses anthrax toxin-encoding genes *cya, lef*, and *pagA* (dark orange) or not (light orange; “Anthrax”); (ii) whether a strain possesses cereulide synthetase-encoding *cesABCD* (dark purple) or not (light purple; “Emetic”); (iii) whether a strain belongs to *panC* Group II (blue) or III (yellow; “*panC”);* (iv) whether a strain was sequenced in this study (dark pink) or not (light pink; “Study”); (v) multi-locus sequence typing (MLST) sequence types (STs) associated with strains sequenced in this study, where applicable (colored), or not (gray; “ST”). MLST lineages discussed in Table 3 are annotated to the right of the heatmap (“MLST Lineage”). MLST lineages with numerical superscripts contain two or more strains sequenced in this study, which were highly similar on a genomic scale; these lineages are depicted in Figure 4. The tree was rooted and time-scaled using LSD2, with branch lengths reported in years.

### A *panC* Group III *B. cereus s.l*. strain with cereulide synthetase-encoding genes belongs to the “high-risk” ST26 lineage

One *panC* Group III *B. cereus s.l*. strain (S66) was assigned to GTDB’s *B. paranthracis* genomospecies and possessed cereulide synthetase-encoding genes (Figure 5 and Table 2). This strain, which had been isolated from processed beef mince from a processing plant in Mpumalanga, was a member of ST26 (Figure 5 and Table 2), the ST to which most cereulide-producing *B. cereus s.l*. strains belong (although it should be noted that ST26 strains may be capable of producing enterotoxins and causing diarrheal illness, regardless of whether they produce cereulide or not) (38, 40, 41). While members of ST26 are comparatively closely related (>99.52 pairwise ANI), WGS was able to distinguish the South African strain sequenced here from closely related ST26 strains (pairwise core SNP distance >913 relative to publicly available genomes; Figure 5 and Table 3).

### A *panC* Group III *B. cereus s.l*. lineage assigned to ST2413 shows evidence of intercontinental dissemination

Two *panC* Group III *B. cereus s.l*. strains sequenced in this study (S59 and S62) were assigned to ST2413 within GTDB’s *B. paranthracis* species (Figure 5 and Table 3). Unlike the ST26 strain, which was also assigned to GTDB’s *B. paranthracis* genomospecies, neither ST2413 strain possessed cereulide synthetase-encoding genes (Figure 5 and Table 2). Both strains were isolated from raw chicken; however, S59 was isolated from a chicken quarter-leg that had been imported from the Netherlands, and S62 from a chicken thigh sold at a retail outlet in Mpumalanga (Figure 4 and Table 1). Notably, these strains were nearly identical on a genomic scale (pairwise core SNP distance = 1, pairwise ANI = 100.0; Figure 5 and Table 3), despite originating from different continents (i.e., Europe and Africa; Figure 4 and Table 1).

### A *panC* Group II *B. cereus s.l*. strain assigned to ST794 is most closely related to a food-associated strain responsible for diarrheal illness in Norway

One *panC* Group II *B. cereus s.l*. strain was sequenced in this study (S57) and was assigned to ST794 within GTDB’s *B. wiedmannii* species (Figure 5 and Table 1). Strain S57 had been isolated from RTE beef biltong sold at a retail outlet in Free State (Table 1). When compared to the two publicly available ST794 genomes, S57 shared > 99.9 ANI with both publicly available genomes and differed by 31 and 100 core SNPs (relative to genomes with NCBI RefSeq Assembly Accessions GCF_900094845.1 and GCF_007671965.1, respectively; Figure 5 and Table 3). Notably, beef biltong-associated strain S57 sequenced here was most closely related to strain NVH 0674-98, a psychrotolerant strain that had been isolated in Norway from mashed swedes, which were reportedly responsible for diarrheal foodborne illness (42). The other ST794 strain, DE0555, was an environmental isolate collected in 2018 from Durham, North Carolina in the United States (NCBI BioSample Accession SAMN11792715).

### A *panC* Group V *B. cereus s.l*. strain from South African mixed-meat wors most closely resembles a plant-associated strain from the United States

One *panC* Group V *B. cereus s.l*. strain was sequenced in this study (S72) and was assigned to ST223 within GTDB’s *B. toyonensis* species (Figure 6 and Tables 1-3). S72 had been isolated in a butchery in Gauteng, from a processed wors composed of a mix of beef, pork, and lamb (Table 1). Strain S72 was most closely related to a publicly available genome, strain AFS092321 (NCBI RefSeq Accession GCF_002552615.1), which had been isolated in 2014 from a tree leaf in North Carolina, United States (NCBI BioSample Accession SAMN07598612) (43); the two strains shared >99.9 ANI and differed by 33 core SNPs (Figure 6 and Table 3).

**Figure 6.**
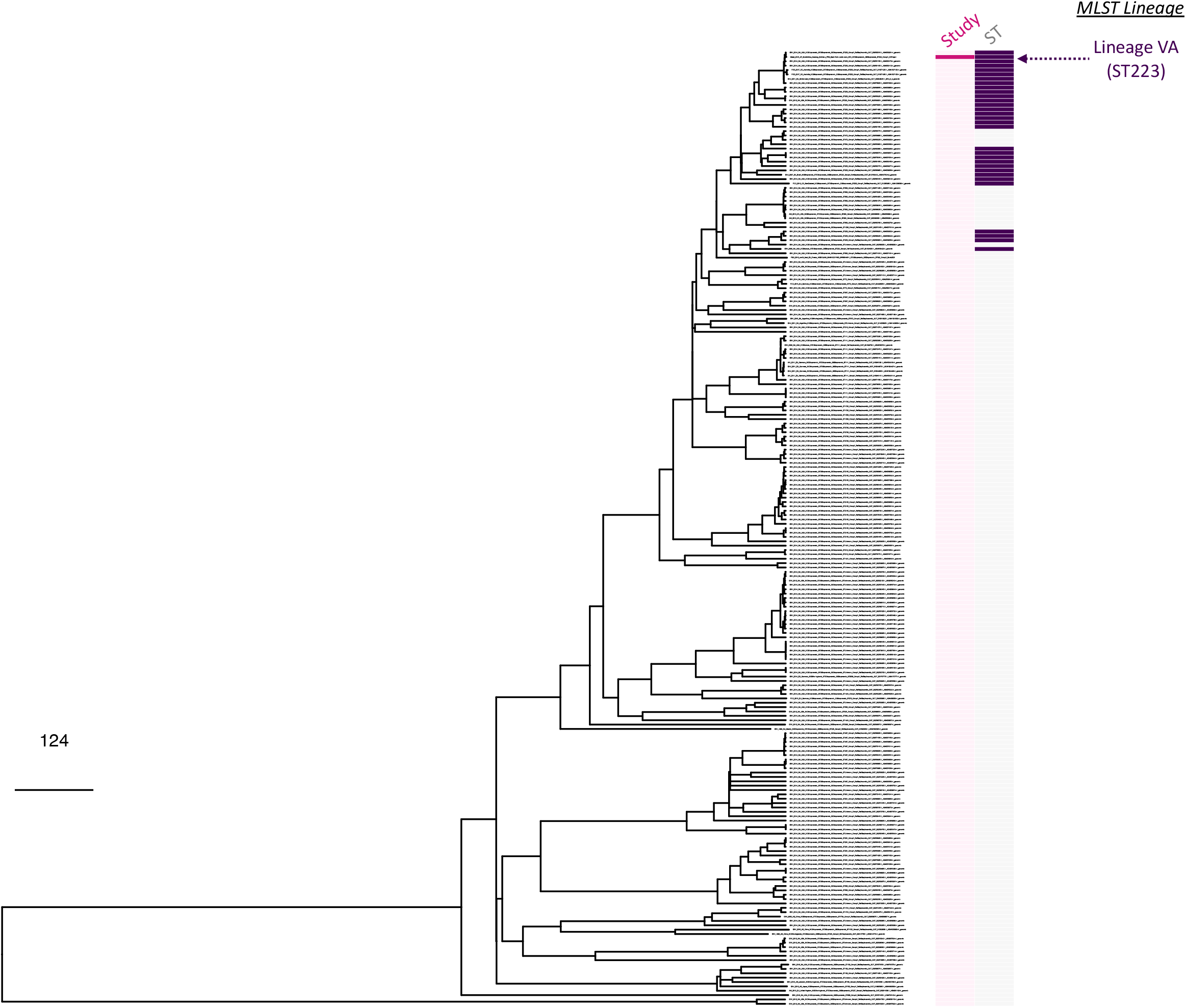
Maximum likelihood phylogeny constructed using core SNPs identified among orthologous gene clusters of 219 *panC* Group V *B. toyonensis* genomes. The phylogeny was rooted using *panC* Group IV *B. cereus* str. ATCC 14579 as an outgroup (NCBI RefSeq Accession GCF_006094295.1; omitted for readability). The heatmap to the right of the phylogeny denotes (i) whether a strain was sequenced in this study (dark pink) or not (light pink; “Study”); (ii) multi-locus sequence typing (MLST) sequence types (STs) associated with strains sequenced in this study, where applicable (colored), or not (gray; “ST”). MLST lineages discussed in Table 3 are annotated to the right of the heatmap. The tree was rooted and time-scaled using LSD2, with branch lengths reported in years.

## DISCUSSION

### *B. cereus s.l*. lineages can be disseminated inter- and intra-nationally via the food supply chain

The movement of commodities (e.g., foods, animals, animal products, agricultural products, consumer products) through inter- and intra-national trade can contribute to the global, regional, and local dissemination of microorganisms, including pathogens (44–48). The international agro-food trade specifically plays an increasingly pivotal role in providing food supplies to communities around the globe but can contribute to the dissemination of foodborne pathogens (47, 48). Consequently, high-resolution technologies such as WGS are being employed increasingly to monitor the spread of pathogens along the food supply chain (49, 50).

Using WGS, we identified six South African *B. cereus s.l*. lineages, which showcased evidence of inter-regional dissemination via the trade and transport of meat products (Figure 4 and Table 3). Notably, one *B. cereus s.l*. lineage showed evidence of intercontinental spread between Europe and Africa: a *B. cereus s.l*. ST2413 strain isolated from raw chicken sold in retail outlets in Mpumalanga, South Africa was identical to a ST2413 strain isolated from chicken imported from the Netherlands. We may hypothesize that the raw chicken sold in Mpumalanga originated from the Netherlands, as the Netherlands was the second-largest exporter of chicken meat products to South Africa in 2014-2016 (i.e., the timeframe in which the strains sequenced here were collected) (51); however, this is merely a hypothesis, as we were unable to confirm this with the retail outlet.

We additionally identified five *B. cereus s.l*. lineages, which showed evidence of inter-provincial spread within South Africa: four *panC* Group IV lineages and one *panC* Group III lineage were each composed of (nearly) identical strains, which were isolated from meat products in two or more South African provinces (Figure 4 and Table 3). Thus, it is likely that strains within each lineage shared a common source; however, a lack of additional metadata and genomes prevents confirmation of this. Overall, these results showcase the utility of WGS for *B. cereus s.l*. source tracking and surveillance, although future studies relying on additional genomes with detailed metadata are needed.

### Nomenclatural frameworks that incorporate both genomic and phenotypic data can improve strain-level *B. cereus s.l*. risk assessment

*B. cereus s.l*. species delineation is notoriously challenging, and numerous *B. cereus s.l*. taxonomic frameworks have been proposed (29). Phenotypic traits historically used for *B. cereus s.l*. species assignment (e.g., motility, colony morphology, ability to cause illness) have long been known to be inconsistent with genome evolution (29, 31, 32). Taxonomies that rely solely on genomic data, however, may lead to incorrect assumptions of a strain’s pathogenic potential, particularly when taxonomic labels have deep roots in medicine or industry (29, 34).

For example, within some ANI-based taxonomic frameworks (e.g., GTDB), the *B. anthracis* genomospecies includes *B. cereus s.l*. strains that, historically, would be referred to as “*B. cereus*” or “Group III *B. cereus*”; these strains possess phenotypic characteristics associated with “*B. cereus*” (e.g., as outlined in the U.S. Food and Drug Administration’s Bacteriological Analytical Manual) (52, 53), and like “*B. cereus*”, they are incapable of causing anthrax (34). These non-anthrax-causing Group III *B. cereus s.l*. strains have been isolated from diverse environments, including food (e.g., milk, egg, whites, spices), consumer products (e.g., baby wipes), and soil, indicating that these organisms are not uncommon in environmental and industrial settings (34). Thus, as WGS becomes more popular in clinical and industrial settings, it is possible that professionals who rely solely on increasingly popular genomic methods for taxonomic delineation (e.g., GTDB, ANI-based comparisons to species type strains) may incorrectly assume that these strains can cause anthrax, due to the “*B. anthracis*” species labels that some taxonomic classification programs produce (29, 34). Here, during routine surveillance of meat products in South Africa, we identified two *panC* Group III *B. cereus s.l*. strains, which did not possess anthrax toxin- or capsule-encoding genes and did not belong to the classic, “clonal” *B. anthracis* lineage associated with anthrax disease (34, 54). These strains would be classified as “*B. cereus*” or “Group III *B. cereus*” using standard microbiological assays (52, 53); however, these strains were assigned to the “*B. anthracis*” genomospecies using GTDB and similar ANI-based methods (Table 1). Referring to these strains as “*B. anthracis*” would be misleading, as they cannot cause anthrax; thus, this potential miscommunication could have disastrous public health and industrial consequences.

We additionally isolated three *B. cereus s.l*. strains from beef and poultry products, which were assigned to the “*B. paranthracis*” genomospecies via GTDB and similar ANI-based methods (Table 1). As noted previously, “*B. paranthracis*” was proposed as a “novel” species in 2017 (55) but was later found to encompass the well-known foodborne pathogen “emetic *B. cereus*” within its genomospecies boundary (29, 37, 38). One of the “*B.paranthracis*” strains isolated here indeed possessed cereulide synthetase-encoding genes and belonged to ST26 (Table 2), the ST that encompasses most *B. cereus s.l*. strains capable of producing emetic toxin (38). This strain thus likely poses a food safety threat and would most likely be referred to as “emetic *B. cereus*” in clinical or industrial settings. Referring to this strain as “*B. paranthracis*” could be misleading to researchers, clinicians, and other professionals who are not well-versed and up to date in *B. cereus s.l*. taxonomy (29, 34, 38). However, not all “*B. paranthracis*” strains are capable of producing emetic toxin: here, two ST2413 strains isolated from poultry were assigned to the “*B. paranthracis*” genomospecies but did not possess cereulide synthetase-encoding genes (Table 2), indicating that these strains cannot cause emetic intoxication. Thus, differentiating potentially emetic from non-emetic strains is critical for informing public health and food safety efforts.

Recently, we proposed a standardized nomenclatural framework for *B. cereus s.l*. (i.e., the 2020 GSB framework), which can utilize genomic, genetic, and/or phenotypic information for taxonomic classification (33, 34). Importantly, the 2020 GSB framework relies on a standardized collection of biovars (i.e., biovar Anthracis, Emeticus, and Thuringiensis), which can be applied to individual *B. cereus s.l*. strains to convey phenotypes of clinical and/or industrial importance (i.e., ability to produce anthrax, emetic, and insecticidal toxins, respectively) (33, 34). Within this framework, the absence of the Anthracis biovar term denotes that *B. cereus s.l*. strains sequenced here *cannot* produce anthrax toxin, while the presence/absence of the Emeticus biovar term differentiates cereulide-producing “*B. paranthracis*” from non-cereulide-producing strains (Table 2). While the 2020 GSB framework provides a standardized set of *B. cereus s.l*. genomospecies (Tables 1 and 2) (33, 34), researchers and other professionals may prefer to use more established names for lineages (e.g., obtained via MLST, *panC* group assignment); biovar terms can thus be appended to *B. cereus s.l*. lineage names (e.g., the ST26 strain sequenced here, which possesses cereulide synthetase, can be referred to as “*B. cereus s.l*. ST26 biovar Emeticus”). Overall, standardized taxonomic frameworks that can incorporate both genomic/genetic and phenotypic information may improve strain-level risk evaluation of *B. cereus s.l*.

### WGS may improve *B. cereus s.l*. genomic surveillance, traceback investigations, and source tracking

WGS has been shown to improve surveillance and source tracking efforts for numerous foodborne pathogens, including *Escherichia coli, Salmonella enterica*, and *Listeria monocytogenes* (50, 56). While the amount of publicly available WGS data derived from members of *B. cereus s.l*. is increasing (34), efforts to sequence the genomes of food-associated *B. cereus s.l*. strains are lagging relative to other foodborne pathogens. Here, we showed that WGS can conceptually be used for *B. cereus s.l*. surveillance and source tracking; however, our study is limited by a lack of publicly available (i) genomic data and (ii) corresponding metadata associated with *B. cereus s.l*. strains. For example, we identified two identical *B. cereus s.l*. ST2413 strains, which were present in both Dutch and South African raw poultry. However, due to a lack of additional publicly available ST2413 *B. cereus s.l*. genomes, we were unable to gain additional insights into exactly where this lineage originated. Thus, future *B. cereus s.l*. surveillance and WGS initiatives in clinical, industrial, and environmental settings are needed to improve *B. cereus s.l*. source tracking and traceback efforts. Furthermore, it is essential that data and metadata obtained in such future initiatives are made publicly available, as international sharing of WGS data can decrease both the amount of time required to solve foodborne outbreaks and the public health burden caused by foodborne pathogens (57). Overall, the proof-of-concept study detailed here highlights the benefits of WGS for *B. cereus s.l*. surveillance and source tracking, even among closely related lineages, and future studies will benefit from increasingly available publicly available WGS data and metadata.

## MATERIALS AND METHODS

### Isolate selection and whole-genome sequencing

A subset of 34 isolates were selected from a total of 79 *B. cereus s.l*. isolates from our previous study (26) using simple random sampling without replacement (58) via random numbers generated in Microsoft Excel. Culturing and genomic DNA extraction was performed as described previously (26), using the High Pure PCR Template preparation kit (Roche, Germany). WGS of selected isolates was performed at the Biotechnology Platform, Agricultural Research Council, Onderstepoort, South Africa. DNA libraries were prepared using TruSeq and Nextera DNA library preparation kits (Illumina, San Diego, CA, USA), followed by sequencing on HiSeq and MiSeq instruments (Illumina, San Diego, CA, USA).

### Data pre-processing and quality control

Paired-end reads associated with each of the 34 isolates were supplied as input to Trimmomatic v0.38 (59), which was used to remove Illumina adapters and leading and trailing low-quality/ambiguous bases (LEADING:3 and TRAILING:3, respectively); reads with average per-base quality scores <15 within a 4 bp sliding window (SLIDINGWINDOW:4:15) were additionally cut, and reads with length < 36 bp were removed. FastQC v0.11.5 (https://www.bioinformatics.babraham.ac.uk/projects/fastqc/) was used to assess the quality of the resulting trimmed paired-end reads.

SKESA v2.4.0 (60) was used to assemble each genome, using the trimmed paired-end reads as input and default settings. SPAdes v3.13.1 (61) was additionally used to assemble each genome in “careful” mode, using trimmed paired-end reads as input. QUAST v4.5 (62) was used to assess the quality of each resulting assembly, and CheckM v1.1.3 (63) was used to evaluate genome contamination/completeness. MultiQC v1.8 (64) was used to assess genome quality in aggregate. Assemblies produced using SKESA were used in all subsequent steps, as they were of higher quality based on metrics produced by QUAST (e.g., N50, number of contigs). Genomes with (i) < 95% completeness (via CheckM), (ii) > 5% contamination (via CheckM), and/or (iii) N50 < 20 Kbp were considered to be of low quality and were excluded (*n* = 4), yielding a preliminary set of 30 genomes used in subsequent analyses.

### *In silico* typing and taxonomic characterization

BTyper3 v3.1.1 (33) was used to characterize each assembled genome (see section “Data pre-processing and quality control” above) using: (i) ANI-based genomospecies, (ii) ANI-based subspecies, and (iii) biovar assignment, using a standardized nomenclatural framework for *B. cereus s.l*. (34) and dependencies FastANI v1.31 (54) and BLAST v2.9.0 (65); (iv) ANI-based pseudo-gene flow unit assignment (33) (also via FastANI); (v) *in silico* seven-gene MLST using the PubMLST *B. cereus* database (accessed 25 October 2020); (vi) *panC* phylogenetic group assignment, using an adjusted eight-group (Group I-VIII) framework (33). All aforementioned analyses were performed using default settings, as well as with virulence factor minimum coverage thresholds lowered to 0% (--virulence_coverage 0) to confirm virulence factor absence. Because BTyper3 uses a conservative approach for Bt toxin gene detection, the command-line implementation of BtToxin_scanner v1.0 (BtToxin_scanner2.pl) was used to identify Bt toxin genes in each genome using default settings (66).

Taxonomic classification of assembled genomes was additionally performed using GTDB Release 05-RS95 (17 July 2020) and GTDB-Tk v. 1.3.0 (i.e., “GTDB R95”), using GTDB-Tk’s “classify_wf” workflow (67–69). Notably, five genomes were assigned to species outside of *B. cereus s.l*. (i.e., three genomes classified as *Escherichia flexneri*, one as *Escherichia dysenteriae*, and one as *Staphylococcus saprophyticus* via GTDB-Tk) and were thus excluded, yielding a final set of 25 *B. cereus s.l*. genomes used in subsequent analyses (Table 1 and Supplemental Table S1).

### Phylogenomic comparison of South African *B. cereus s.l*. genomes to *B. cereus s.l*. species type strains

Prokka v1.14.6 (70) was used to annotate each of the 25 *B. cereus s.l*. genomes sequenced in this study. Protein coding sequences derived from the type strain genomes of each of the 23 validly published and effective *B. cereus s.l*. species (accessed 28 August 2021) were downloaded from NCBI’s RefSeq Assembly database (see Table 1 of Méndez Acevedo, et al. for all type strain accessions) (71). OrthoFinder v2.5.2 (72, 73) was used to identify orthologues among protein coding sequences associated with all 47 genomes (25 *B. cereus s.l*. genomes sequenced in this study, plus 23 *B. cereus s.l*. species type strain genomes), using MAFFT v7.475 (74, 75) for sequence alignment and RAxML-NG v1.0.2 (76) for phylogeny construction.

The resulting amino acid sequence alignment was supplied as input to IQ-TREE v1.5.4 (77), which was used to construct a maximum likelihood (ML) phylogeny, using 1,000 replicates of the ultrafast bootstrap approximation (78), plus the optimal amino acid substitution model selected using ModelFinder (i.e., the general matrix model with empirical amino acid frequencies and the FreeRate model with six categories; JTT+F+R6) (79–82). The resulting phylogeny was rooted using effective species “*B. manliponensis*” (i.e., the most distant recognized member of *B. cereus s.l.*) (83) and annotated using the bactaxR package (34) in R v4.1.2 (84).

### Acquisition of publicly available *B. cereus s.l*. genomes

All assembled genomes submitted to the National Center for Biotechnology Information (NCBI) RefSeq database (85, 86) as one of 23 validly published or effective *B. cereus s.l*. species (i.e., *albus, anthracis, “bingmayongensis”, cereus, “clarus”, cytotoxicus, fungorum, “gaemokensis”, luti*, “*manliponensis”, mobilis, mycoides, nitratireducens, pacificus, paramycoides, paranthracis, proteolyticus, pseudomycoides, thuringiensis, toyonensis, tropicus, weihenstephanensis, wiedmannii*) (1, 29, 55, 71, 83, 87–99) were downloaded (*n* = 2,733; accessed 20 March 2021). QUAST and CheckM were used to assess the quality of each assembled genome (see section “Data pre-processing and quality control” above), and BTyper3 (using default settings) and GTDB-Tk were used for typing and/or taxonomic assignment as described above (see section “*In silico* typing and taxonomic characterization”). The rentrez package (v1.2.3) was used to download metadata associated with each genome’s BioSample in R v3.6.1 (84, 100, 101). Publicly available genomes meeting all of the following quality thresholds were used in subsequent analyses (*n* = 2,664; Supplemental Table S2): (i) > 95 % completeness (via CheckM), (ii) < 5% contamination (via CheckM), (iii) N50 > 20 Kbp (via QUAST), (iv) composed of < 1,000 contigs (via QUAST).

### Acquisition of genomes from a study of *B. thuringiensis* outbreaks

All sequencing reads associated with isolates from a previous study of outbreaks caused by *B. thuringiensis* in France (102) were downloaded from NCBI’s Sequence Read Archive (SRA) database using the SRA Toolkit v 2.8.2 (103, 104). Genomic data for all 171 isolates were pre-processed, assembled, and taxonomically classified as described above, with genomes assembled using SKESA used in subsequent steps (see sections “Data pre-processing and quality control” and “*In silico* typing and taxonomic characterization” above). Four genomes did not meet the quality thresholds employed in this study (see section “Acquisition of publicly available *B. cereus s.l*. genomes” above) and were thus excluded, yielding 167 genomes from the study, which were used in subsequent analyses (Supplemental Table S3).

### Acquisition of genomes derived from strains isolated in conjunction with a previous outbreak caused by emetic *B. cereus s.l*

The genomes of 33 *B. cereus s.l*. strains isolated in conjunction with a 2016 emetic outbreak in New York State (United States) were downloaded, preprocessed, and assembled as described previously (37). The quality of each of the 33 genomes was assessed as described above (see section “Data pre-processing and quality control” above), and all genomes underwent taxonomic classification and typing as described above (see section and “*In silico* typing and taxonomic characterization” above). Two genomes did not meet the quality thresholds employed in this study (see section “Acquisition of publicly available *B. cereus s.l*. genomes” above) and were thus excluded, yielding 31 genomes from the study, which were used in subsequent analyses (Supplemental Table S4).

### Within-group phylogeny construction

The 25 *B. cereus s.l*. strains sequenced here spanned four major phylogenetic groups, based on their *panC* sequence (i.e., *panC* Groups II, III, IV, and V; Table 1). Thus, phylogenies were constructed using all genomes assigned to each of the following major lineages: (i)*panC* Group IV (Figure 3), (ii)*panC* Groups II and III (Figure 5), and (iii)*panC* Group V (Figure 6), which are equivalent to the (i) *B. cereus s.s.*, (ii) *B. mosaicus*, and (iii) *B. toyonensis* genomospecies within the 2020 GSB taxonomic framework (33), respectively (*panC* Group II and III genomes were aggregated, due to the fact that these lineages are closely related and polyphyletic; Figure 5).

For each of the three major lineages, Prokka was used to annotate each genome; the resulting GFF files associated with each genome were supplied as input to Panaroo v1.2.8 (105), which was used to partition genes into core- and pan-genome orthologous gene clusters, using the following parameters (all other parameters were set to their default values): (i) “strict” mode (--clean-mode strict), (ii) core genome alignment using MAFFT (-a core --aligner mafft), (iii) a core genome sample threshold of 95% (--core_threshold 0.95). The resulting core genome (nucleotide) alignment was queried using snp-sites v2.5.1 (106), which was used to identify (i) core SNPs and (ii) constant sites among all genomes in the major lineage. The resulting core SNP alignment was supplied as input to IQ-TREE v1.5.4, which was used to construct a ML phylogeny using the General Time-Reversible (GTR) nucleotide substitution model (107), one thousand replicates of the ultrafast bootstrap approximation (78), and an ascertainment bias correction obtained using constant sites output by snp-sites.

For each of the three major lineages, all aforementioned steps were repeated, with the addition of an outgroup genome: for the *panC* Group IV phylogeny, *panC* Group III *B. anthracis* str. Ames Ancestor was used as an outgroup (NCBI RefSeq Accession GCF_000008445.1); for the *panC* Groups II/III and *panC* Group V phylogenies,*panC* Group IV *B. cereus* str. ATCC 14579 was used as an outgroup (NCBI RefSeq Accession GCF_006094295.1). Additionally, only genomes with detailed metadata (i.e., a reported year of isolation, isolation source, and geographic location) were included in this analysis (Supplemental Tables S1-S4). Each of the three resulting phylogenies were rooted and scaled using LSD2 v1.4.2.2 (108) and the following parameters: (i) tip dates corresponding to the year of isolation associated with each genome; (ii) constrained mode (-c), with the root estimated using constraints on all branches (-r as); (iii) variances calculated using input branch lengths (-v 1); (iv) 1,000 samples for calculating confidence intervals for estimated dates (-f 1000); (v) a sequence length of 5,500,000 (-s 5500000); (vi) rooting along the outgroup genome (-g -k). The resulting phylogenies were annotated using the bactaxR package in R.

### Delineation of MLST lineages and identification of closely related and “identical” genomes

FastANI v1.31 was used to calculate ANI values between each of the 25 *B. cereus s.l*. genomes sequenced in this study (i.e., as a query genome), and all genomes assigned to the *panC* Group of the query genome (*panC* Groups II and III were aggregated); genomes were then grouped into lineages based on STs assigned using seven-gene MLST (see section “*In silico* typing and taxonomic characterization” above). For each of the resulting MLST lineages, FastANI was used to calculate pairwise ANI values between all genomes within the MLST lineage (Table 3).

For each MLST lineage, Snippy v4.6.0 (https://github.com/tseemann/snippy) was used to identify core SNPs among all genomes assigned to the respective MLST lineage, using (i) a genome sequenced in this study as a reference genome (Table 3); (ii) paired-end reads associated with each genome as input (for the genomes sequenced in this study, as well as the genomes from the Bonis, et al. study and the New York State outbreak study) (37, 102) and/or assembled genomes as input (for NCBI genomes); (iii) default settings. For MLST lineages with more than four genomes (i.e., ST24, ST26, ST177, ST223, ST1578), Gubbins v3.1.3 (109) was used to remove recombination, and core SNPs were identified within the resulting filtered alignment using snp-sites. For all MLST lineages, pairwise core SNP distances were calculated within the MLST lineage (i) among all genomes, (ii) among genomes sequenced in this study, and (iii) between genomes sequenced in this study and publicly available genomes (Table 3), using the dist.gene function in the ape package (110, 111) in R.

### Data availability

Paired-end Illumina reads associated with the 25 *B. cereus s.l*. isolates sequenced in this study have been deposited in NCBI’s SRA database under BioProject Accession PRJNA798224. Metadata and quality information for all genomes queried in this study are available in Supplemental Table S1 (the 25 isolates sequenced in this study) and Supplemental Tables S2-S4 (all publicly available genomes).

## ACKNOWLEDGMENTS

The following organisations are acknowledged for their contributions:

- Gauteng Department of Agriculture and Rural Development (GDRAD) for project funding and the use of data for this study.
- The authors are grateful to the Agricultural Research Council: Onderstepoort Veterinary Research for providing all research facilities.
- Our collaborator at EMBL
- Figures 1 and 4 were created with BioRender.com.

## AUTHOR CONTRIBUTIONS

LC designed and carried out all computational analyses. IM and LC conceptualized the study. RP and IM sourced the funding for the project. RP supervised the sequencing of the isolates, while MM and AA performed all culturing work and DNA extractions. LC and IM co-wrote the manuscript, with input from all authors. All authors contributed to the article and approved the submitted version.

